# Symmetry processing in the macaque visual cortex

**DOI:** 10.1101/2021.03.13.435181

**Authors:** Pauline Audurier, Yseult Héjja-Brichard, Vanessa De Castro, Peter J. Kohler, Anthony M. Norcia, Jean-Baptiste Durand, Benoit R. Cottereau

## Abstract

Symmetry is a highly salient feature of the natural world that is perceived by many species of the animal kingdom and that impacts a large array of behaviours such as partner selection or food choice. In humans, the cerebral areas processing symmetry are now well identified from neuroimaging measurements. However, we currently lack an animal model to explore the underlying neural mechanisms. Macaque is a potentially good candidate, but a previous comparative study (1) found that functional magnetic resonance imaging (fMRI) responses to mirror symmetry in this species were substantially weaker than those observed in humans under similar experimental conditions. Here, we re-examined symmetry processing in macaques from a broader perspective, using both rotation (experiment 1) and reflection (experiment 2) symmetry. Our experimental design was directly derived from that of a previous human fMRI study (2), in order to facilitate the comparison between the two primate species. Highly consistent responses to symmetry were found in a large network of areas (notably V3, V3A, V4, V4A and PITd), in line with what has been observed in humans. Within this network, response properties in areas V3 and V4 (notably their dependency on the rotation symmetry order) were strikingly similar to those observed in their human counterparts. Our results suggest that the cortical networks that process symmetry in humans and macaques are much more similar than previously reported and point toward macaque as a relevant model for understanding symmetry processing.

**Significance statement:** Symmetry processing is an important aspect of human visual perception. We currently lack an animal model for characterizing the neural mechanisms that underlie it at the microscopic scale. Here, we use fMRI measurements in macaques to demonstrate that the cortical responses to symmetry in this species are comparable to those observed in humans under similar experimental conditions to a much higher extent than previously documented. Our results call for a re-examination of the relevance of the macaque model for symmetry processing in humans and open the door to an exploration of the underlying neural mechanisms at the single-cell level, notably in V3, an area often neglected in most current models of visual processing.

## Introduction

As written by the physicist and essayist Alan Lightman: “The deep question is: Why does nature embody so much symmetry? We do not know the full answer to this question” (3). Symmetry is indeed prevalent in natural scenes and a large variety of animals like birds (4), fishes (5) or even insects (6) are able to detect it, notably as a marker of phenotypic and/or genotypic quality in potential partners (7). In humans, symmetry perception has been well documented by psychologists (see ref. 8 for a review) and could underlie some aspects of our aesthetic judgments (9) as demonstrated by its omnipresence in art, craft, and architecture. From the perspective of neuroimaging, the cortical networks involved in aspects of symmetry processing are beginning to be understood. They include extrastriate visual areas like V3 and V4 as well as higher-level regions in both the dorsal (e.g. area V3A) and ventral (e.g. areas VO1, VO2 or LO) pathways (1,2,10,11). While these studies have provided important information regarding where in the cortical processing hierarchy symmetry is represented, they cannot address questions of the underlying cellular mechanisms involved in symmetry processing. In this context, macaques could constitute a promising animal model because they perceive symmetry (12) and it is established that the functional organisation of their visual system is substantially similar to that of Humans (13,14). To assess the utility of a macaque model for symmetry processing in humans, it is important to first determine whether the cortical areas with significant responses to symmetry are analogous in these two primate species.

So far, only one study has explored symmetry processing in humans and monkeys using a comparative approach. In this study, individuals of both species were exposed to random dot patterns with (or without) reflection symmetry (1). Extensive measurements using standard experimental conditions at 3 Teslas failed to reveal significant responses to reflection symmetry in macaque. Only by using contrast agents or high-field functional magnetic resonance imaging (fMRI) at 7 Teslas could the authors detect symmetry-related activations in this species. These activations were only observed within areas V4d and V3A, suggesting a much more restricted cortical network than the one observed in humans using the same protocol. The fact that symmetry was found to evoke weak cortical responses in macaque possibly discouraged further explorations (notably in electrophysiology) in that model. To our knowledge, there has been no study on symmetry processing in the non-human primate brain since this single publication, more than 15 years ago.

Here, we re-examined the fMRI responses to symmetry in macaque from a broader perspective. We used both rotation and reflection symmetry and also manipulated the amount of symmetry in our stimuli to determine whether it modulates cortical activations, as observed in humans (1,2). To facilitate the comparison between the two primate species, our experimental design was directly derived from that of a previous human study using regular textures (wallpaper patterns, 15).

Our results clearly establish that macaque brains process symmetry using a much broader cortical network than previously documented, and that the areas involved are closely related to those observed in humans. Thus, the present study calls for a re-examination of the relevance of the macaque model for symmetry processing in humans and it opens the door to a characterization of the underlying neural mechanisms at the single-cell level, notably in area V3.

## Results

The aim of this study was to characterize the cortical areas that process different forms of symmetry in macaque. We recorded fMRI activations to rotation (experiment 1) and reflection (experiment 2) symmetries in two awake behaving animals (M01 and M02) involved in a passive fixation task. Our fMRI contrasts were based on control stimuli defined from a phase scrambling in the Fourier domain (figure 1-A, see also ref 2). This operation modifies the global spatial content (and notably the symmetry properties) without affecting the amplitude spectrum (i.e. it preserves the local properties of the stimuli; see the ‘Materials and Methods’ section). We used a blocked-design during which periods of visual stimulation (with either symmetric stimuli or their phase-scrambled controls) were interleaved with periods of fixation on a gray screen.

**Figure 1:**
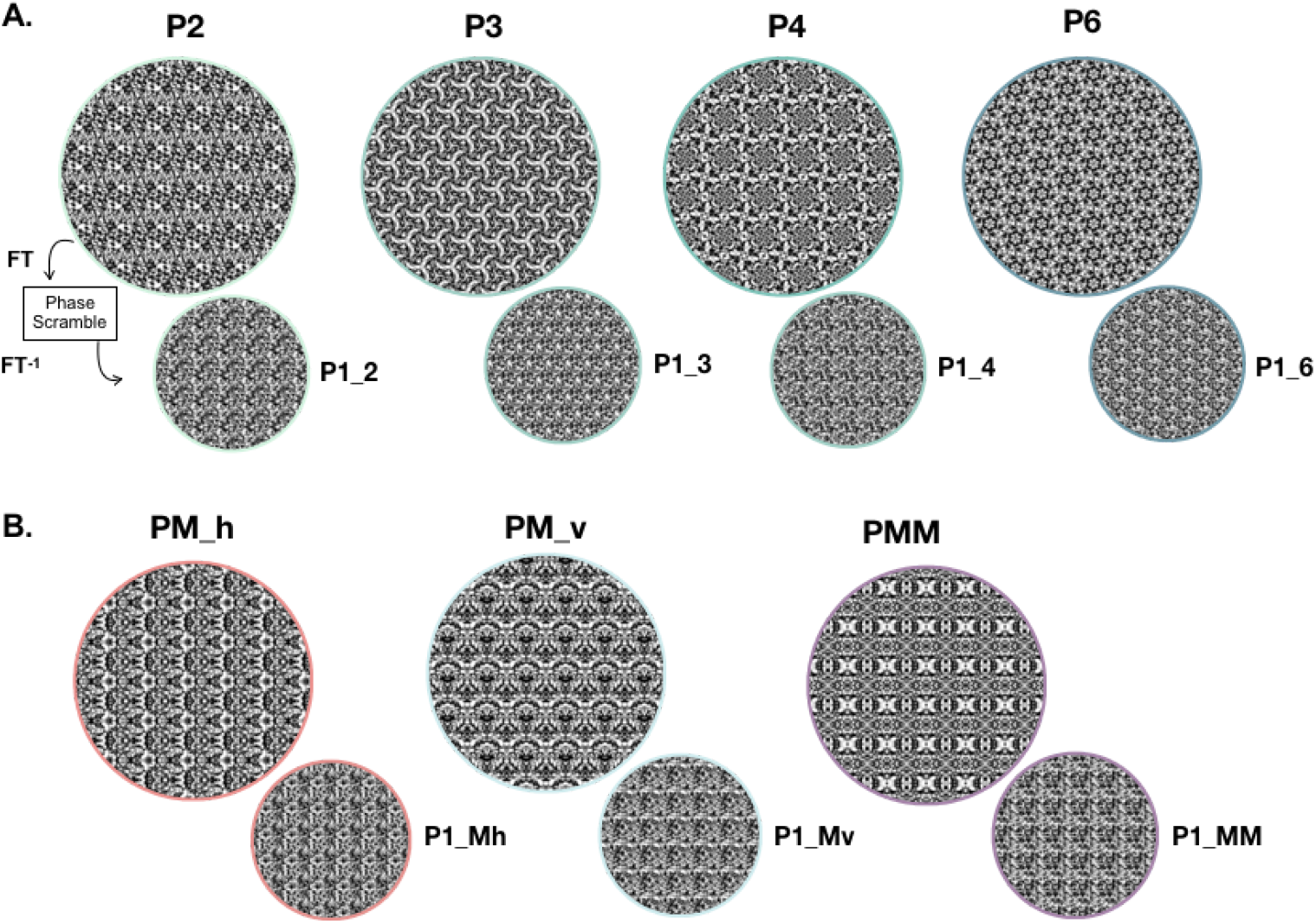
Visual stimuli. A) Exemplar images with rotation symmetry (experiment 1) of orders 2 (P2), 3 (P3), 4 (P4) and 6 (P6). For each of these examples, a control condition (P1) was defined by scrambling the phases of the stimulus in the Fourier domain. Each stimulus and its control have the same power spectrum and therefore share the same low-level properties. B) Exemplar images with reflection symmetry (experiment 2) and their respective controls. Reflection symmetries were based on horizontal axes (PM_h), vertical axes (PM_v) or a combination of both (PMM).

### Experiment 1

#### Responses to rotation symmetry

The experimental design was similar to that used in a previous human fMRI study (2), allowing a direct comparison of the cortical networks processing rotation symmetry in the two primate species. We first examined the differences in blood oxygen level-dependent (BOLD) responses evoked by the rotation symmetry stimuli (all orders pooled together) and by their respective phase-scrambled controls (see figure 1-A). Figure 2-A presents the corresponding statistical parametric maps (t-scores) projected on dorsal and lateral views of inflated reconstructions of the left and right cortical hemispheres of our two monkeys (M01 and M02). Projections on ventral and medial views, for which we did not observe significant activations, are provided in supplementary figure 1-A. Hot colors (orange to yellow) indicate significantly stronger BOLD activation for symmetry (t-score > 3, p-value < 10−3 uncorrected). Very similar activation patterns were observed in the visual cortices of our two animals (activation overlaps on the F99 macaque template are provided in supplementary figure 2-A). A set of retinotopic areas was independently delineated for each monkey (see ref 16,17) and overlaid on the activation map. This retinotopic parcellation reveals sensitivity to rotation symmetry in visual areas V2, V3, V4 and V3A. This result was confirmed by ROI-based analyses in all the retinotopically-defined areas, including those of the middle temporal (MT) and posterior intra-parietal (PIP) clusters (see figure 2-B, values in the satellite areas of the MT and PIP clusters are provided in supplementary figure 3-A). T-score values in V2, V3, V4 and V3A were consistently greater than three for both animals and in both hemispheres. Stronger responses to symmetry were also observed within the infero-temporal gyrus, anterior to area V4 (see the cyan circles), as described in detail below.

**Figure 2:**
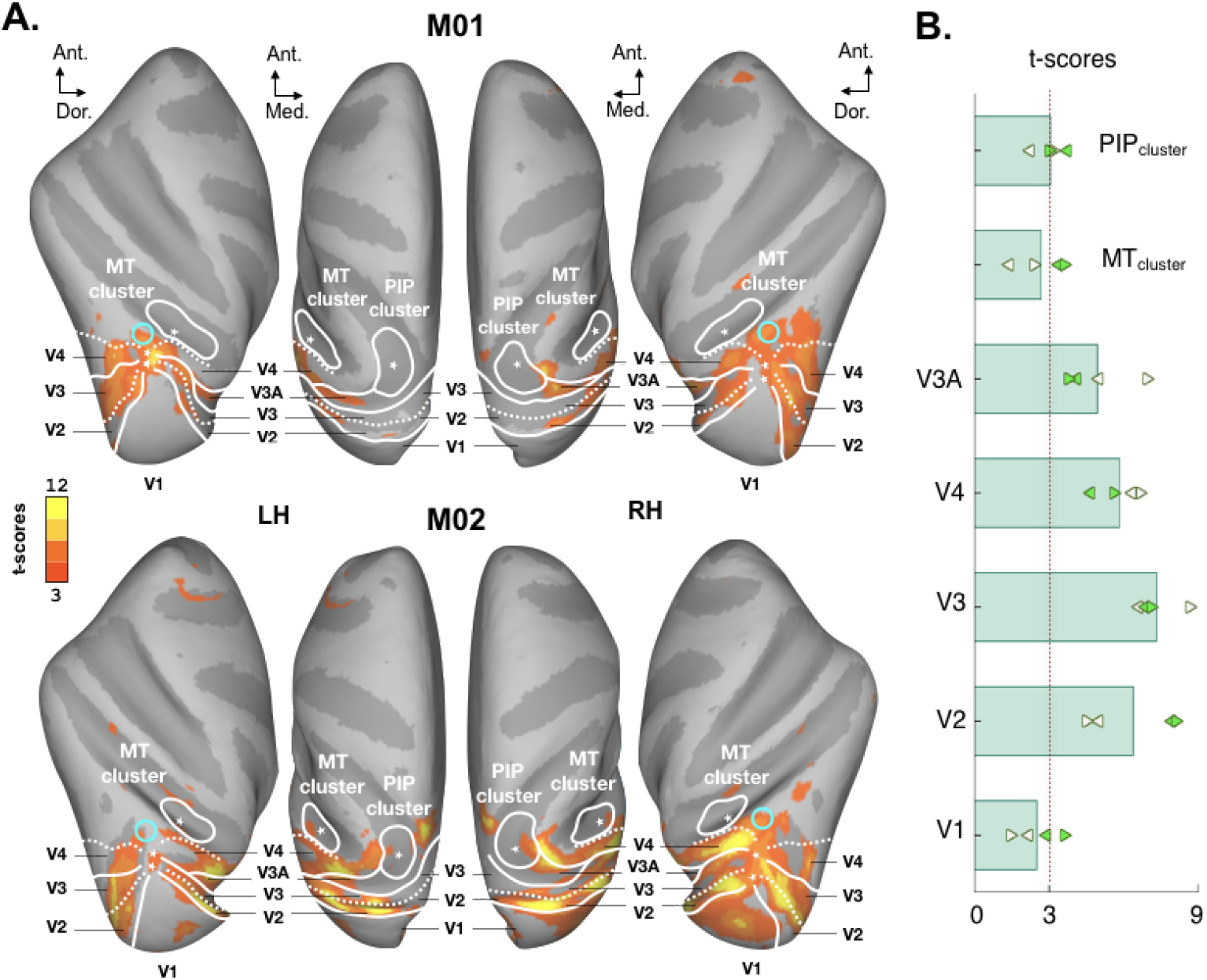
Comparison between BOLD responses to the rotation symmetry (all orders) and to the control conditions. A) Whole brain univariate statistical maps. Activations that were stronger for symmetry were projected on the individual cortical surface (dorsal and lateral views) of M01 (upper panel) and M02 (lower panel). Data were thresholded at p-value < 10−3 (uncorrected). Limits between visual areas (V1, V2, V3, V4 and V3A) obtained from an independent retinotopic mapping protocol are marked by solid (representation of a vertical meridian of the visual field) and dotted (representation of a horizontal meridian) white lines. The MT and PIP clusters (also defined from the retinotopic mapping protocol) are provided by solid white contours. Foveal confluences are marked by stars. Cyan circles show activations beyond retinotopic areas. Ant.: anterior, Dor.: dorsal. Med.: medial. MT: middle temporal, PIP: posterior intra-parietal, LH: left hemisphere, RH: right hemisphere. ROI-based statistics. T-scores for the symmetry versus control conditions within areas V1, V2, V3, V4 and V3A and within the MT and PIP clusters. These values were averaged across the two monkeys and are given by the green bars. Left and right arrows provide t-scores in the left and right hemisphere for M01 (white arrows) and M02 (green arrows). The red dotted line corresponds to the threshold (t-score = 3) used in panel A.

**Figure 3:**
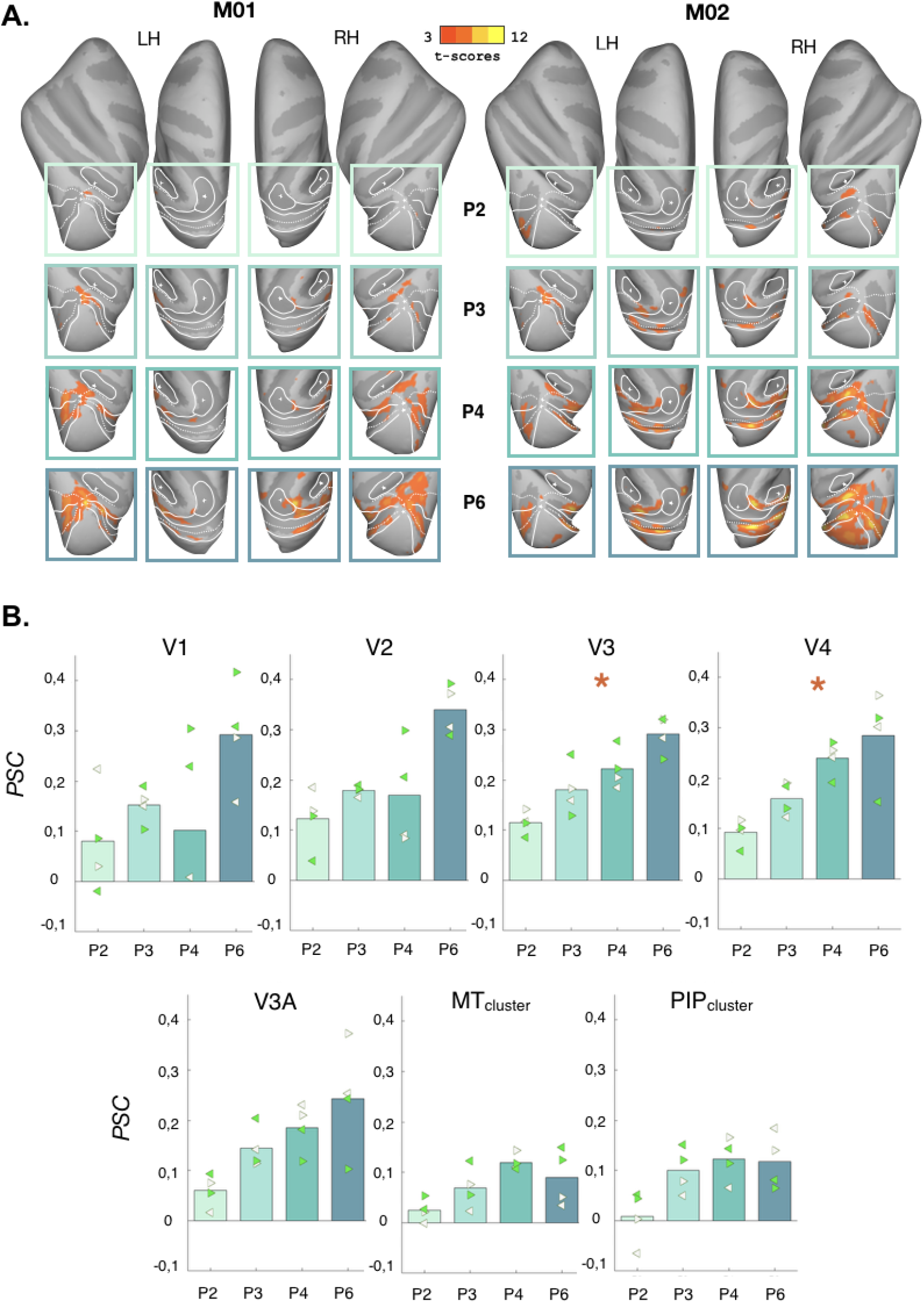
Effects of rotation symmetry order on BOLD responses. A) Whole brain univariate statistical maps. Activations that were stronger for each symmetry order (P2, P3, P4 and P6) with respect to their control conditions are shown in the different boxes for M01 (leftward columns) and M02 (rightward columns). Data were thresholded at p-value < 10−3 (uncorrected). See figure 2-A for more details. B) Percentages of signal changes (PSCs) between the responses to the rotation symmetry conditions (P2, P3, P4 and P6) and those to their respective controls. Data are shown in retinotopic areas (V1, V2, V3, V3A and V4) and in the MT and PIP clusters. Left and right arrows provide values in the left and right hemisphere for M01 (white arrows) and M02 (green arrows). Areas marked with a star (‘*’) are those for which we found a significant linear relationship between PSCs and symmetry order in both the two animals.

#### Effects of rotation symmetry order

In humans, some visual areas exhibit BOLD responses proportional to the rotation symmetry order (2). In order to test whether such an effect also exists in macaque, we first examined the whole brain statistical maps corresponding to the difference in BOLD signal between the four wallpaper groups (P2, P3, P4 and P6) and their respective controls (see figure 3-A). There is a clear tendency for activations to increase with symmetry order in the left and right hemispheres of both monkeys. Next, we computed the corresponding percentages of signal change (PSC) for all retinotopic ROIs (including the MT and PIP clusters). These values are shown in figure 3-B (values in the areas constituting the MT and PIP clusters are provided in supplementary figure 4-A, individual data for M01 and M02 are shown in supplementary figure 5). We also performed linear regressions between these PSCs and the symmetry orders, independently for each animal and for each ROI (averaged across hemispheres). We found significant linear relationships in both monkeys for areas V3 (t-score = 3.2, p-value = 0.0022 in M01 and t-score = 2.513, p-value = 0.0136 in M02) and V4 (t-score = 4.193, p-value = 0.0001 in M01 and t-score = 2.472, p-value = 0.0152 in M02). The corresponding equations and the variances explained by these linear models are provided in supplementary table 1. We also found significant linear relationships in V2 for M02 and in V3A for M01.

**Figure 4:**
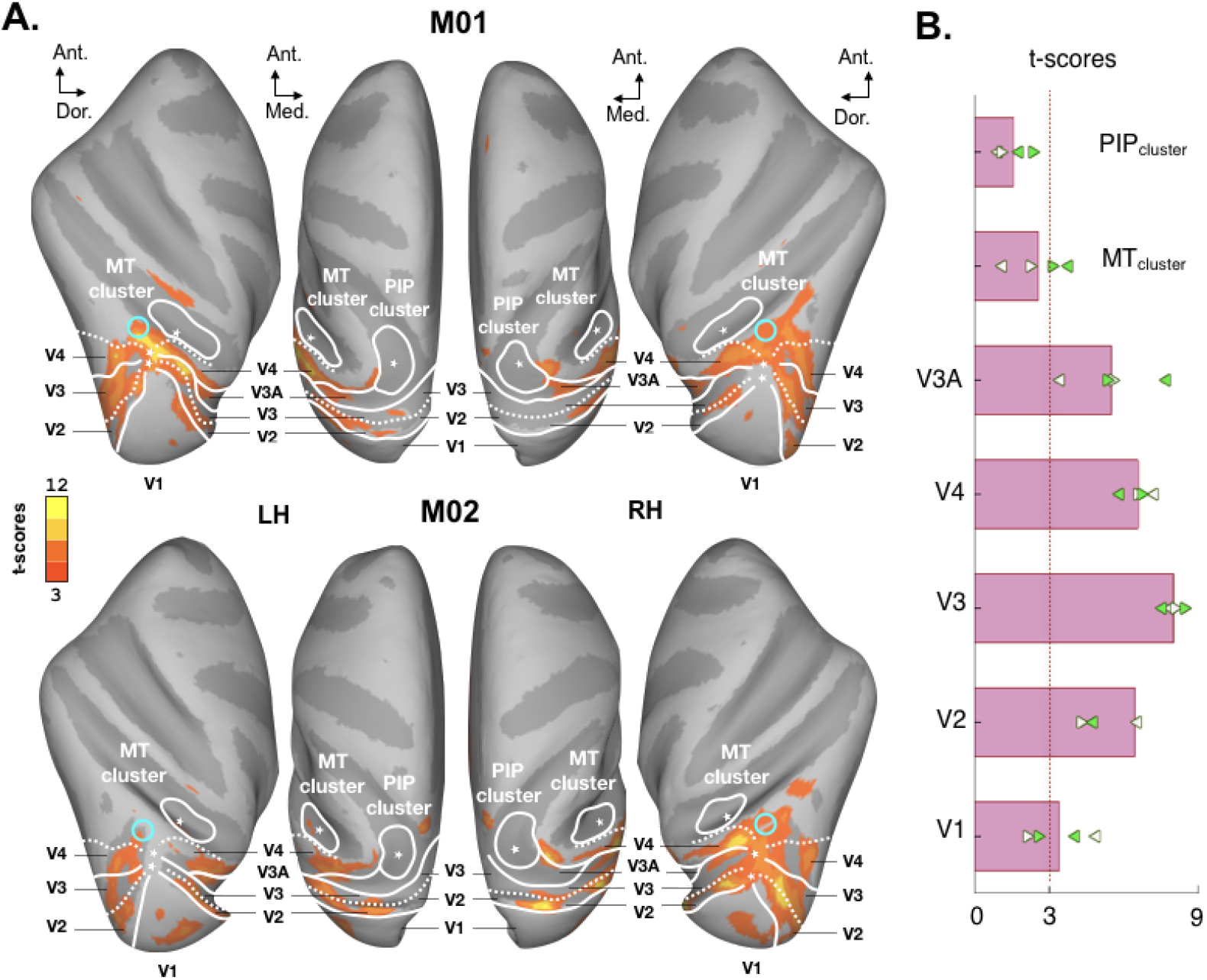
Comparison between responses to reflection symmetry (all conditions) and to the control conditions. A) Whole brain univariate statistical maps. See figure 2-A for more details. B) ROI-based statistics. See figure 2-B for more details.

**Figure 5:**
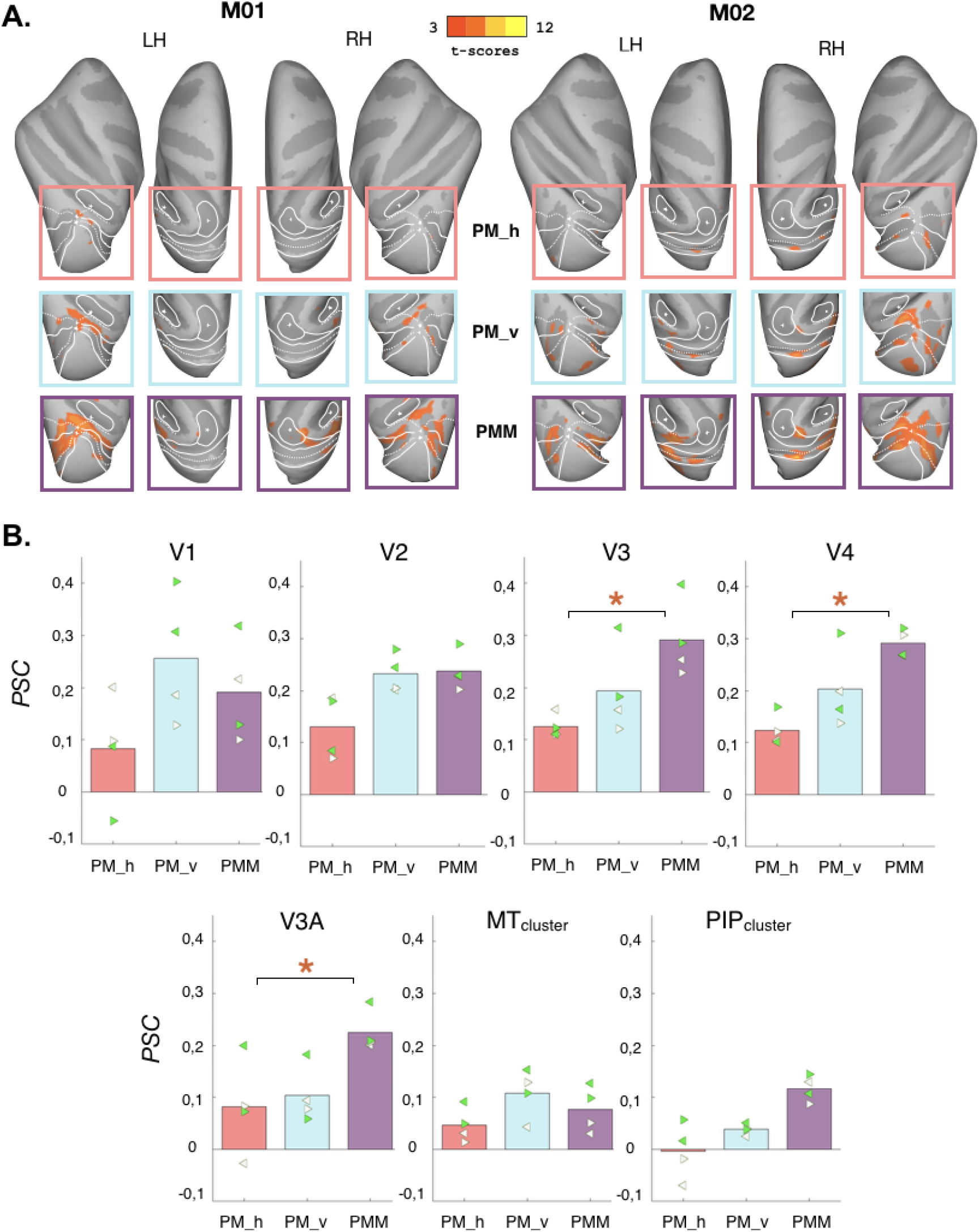
Effects of the different reflection symmetry conditions on BOLD responses. A) Whole brain univariate statistical maps. Responses that were stronger for each symmetry order (PM_h, PM_v and PMM) with respect to their control conditions are shown in the different boxes for M01 (leftward columns) and M02 (rightward columns). See figures 2-A and 3-A for more details. B) Percentages of signal changes (PSCs) obtained for each of the reflection symmetry conditions (PM_h, PM_v and PMM) versus their respective controls. See figure 3-B for more details. Stars indicate ROIs for which the non-parametric permutation tests found that PMM elicited significantly stronger responses than PM_h in both of the two animals.

### Experiment 2

#### Responses to reflection symmetry

Next, we characterized fMRI responses to images containing different axes and amounts of reflection symmetry. Stimuli were wallpaper groups that included reflection symmetry with horizontally oriented axes (PM_h), vertically oriented axes (PM_v) or both (PMM) (see figure 1-B). We first examined BOLD signal differences between the responses evoked by the symmetry stimuli (all conditions pooled together) and their P1 controls. Figure 4-A presents the corresponding statistical parametric maps (t-scores) projected on dorsal and lateral views for M01 and M02 (projections on ventral and medial views are provided in supplementary figure 1-B). Response patterns are very similar between the two animals (see the overlapping map in supplementary figure 2-B) and also match very closely those observed for rotation symmetry (overlaps between the two types of symmetry are provided in supplementary figure 2-C). As was the case for rotation, the most responsive areas are V2, V3, V4 and V3A with t-scores greater than 3 in the two hemispheres of the two monkeys (see the ROI-based statistics in figure 4-B, t-scores in the satellite areas of the MT and PIP clusters are provided in supplementary figure 3-B). Stronger responses to symmetry were also observed within the infero-temporal gyrus, anterior to area V4 (see the cyan circles).

#### Effects of the axes of symmetry

In the first experiment, we found that some visual areas exhibit BOLD responses proportional to the rotation symmetry order. For reflection symmetry, responses could therefore be more pronounced for conditions with more symmetry axes (i.e. in the PMM condition). In order to test this hypothesis, we examined the statistical maps corresponding to the difference in BOLD signal between each reflection symmetry condition (PM_h, PM_v and PMM) and their respective controls (see figure 5-A, only dorsal and lateral views are shown here, ventral and medial views are provided in supplementary figure 1-B). Responses are generally more pronounced for the PMM condition. The corresponding PSCs in the retinotopic ROIs and in the MT and PIP clusters are provided in figure 5-B (values within the areas of the MT and PIP clusters are provided in supplementary figure 4-B, individual data for M01 and M02 are shown in supplementary figure 6).

**Figure 6.**
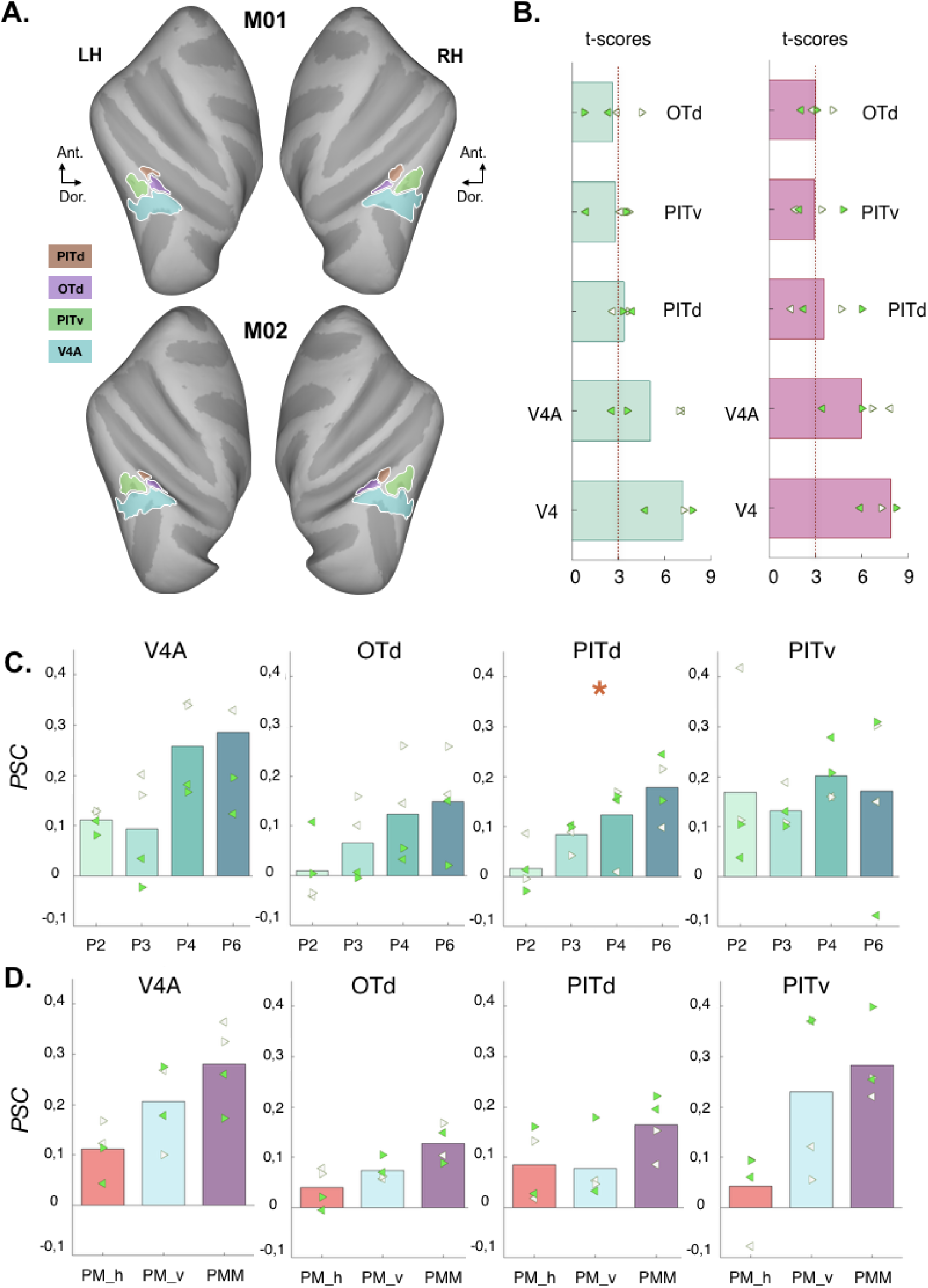
A) Areas V4A, OTd, PITv and PITd defined from the probabilistic atlas of Janssens and Vanduffel and shown on lateral views of inflated reconstructions of the left and right cortical hemispheres of our two monkeys (M01 and M02). B) T-scores for the symmetry versus control conditions within these areas. Values for rotation (experiment 1) and reflection (experiment 2) symmetries are respectively shown on the left and on the right. Values were averaged across the two monkeys. Left and right arrows provide t-scores in the left and right hemisphere for M01 (white arrows) and M02 (green arrows). C) Percentages of signal changes (PSCs) between the responses to the rotation symmetry conditions (P2, P3, P4 and P6) and those to their respective controls. Left and right arrows provide values in the left and right hemisphere for M01 (white arrows) and M02 (green arrows). Areas marked with a star (‘*’) are those for which we found a significant linear relationship between PSCs and symmetry order in both the two animals. D) Percentages of signal changes (PSCs) obtained for each of the reflection symmetry conditions (PM_h, PM_v and PMM) versus their respective controls. See panel-C for more details.

Non-parametric permutation tests demonstrated that stimuli containing both horizontal and vertical symmetries (PMM) elicited significantly stronger responses than those with only horizontal symmetries (PM_h) for both monkeys in areas V3 (p-value = 0.0021 in M01 and p-value = 0.0043 in M02), V4 (p-value = 0.0035 in M01 and p-value = 0.0028 in M02) and V3A (p-value = 0.0034 in M01 and p-value = 0.0022 in M02). We also found that PMM stimuli evoked significantly stronger responses than stimuli containing vertical symmetries (PM_v) in areas V3A for M02 and in V3 and V4 for M01.

#### Symmetry responses beyond V4 in atlas-based ROIs

For both rotation and reflection symmetries, significant responses were observed beyond V4, within the infero-temporal gyrus (see the cyan circles in figures 2-A and 4-A). To further characterize these responses, we performed ROI-based analyses in the V4A area, in the dorsal occipitotemporal area (OTd) and in the ventral and dorsal posterior inferotemporal areas (PITv and PITd) defined from the probabilistic atlas of Janssens and Vanduffel (see the materials and methods section and ref. 37). The definition of these ROIs on the inflated reconstructions of the left and right cortical hemispheres of our two monkeys are shown in figure 6-A (lateral views). T-scores for the two types of symmetry are given in figure 6-B (t-scores in area V4 defined from the same atlas are also provided for comparison with values in retinotopically defined V4). As for the retinotopically-defined ROIs (see figures 2-B and 4-B), there is a very good correspondence between the statistical values obtained for rotation and reflection symmetry. Beyond V4, the most responsive area is V4A with t-scores greater than 3 for both experiments. By comparison, symmetry responses in areas OTd, PITv and PITd are only moderate (t-scores around 3). However, as for areas V3 and V4 (see figure 3-B), we found a significant linear relationship between PSC and rotation symmetry order for area PITd for both animals (t-score = 4,639, p-value = 0,035 in M01 and t-score = 8,02, p-value = 0,005 in M02, see figure 6-C and supplementary table 1 for the associated equation and variance explained). Significant linear relationships were also found in area OTd for M01. We did not find significant difference between responses to the different reflection symmetry conditions in the same area for both monkeys (figure 6-D), even though permutation tests showed that responses to PMM were stronger than those to PM_h in area V4A and PITv for M01 and PITd for M02.

## Discussion

The aim of this study was to characterize the areas in macaque cortex that process symmetry and to determine whether these areas have potential counterparts among the set of areas observed in humans (1,2,10,11). We recorded fMRI activations to rotation (experiment 1) and reflection (experiment 2) symmetries in two awake behaving animals involved in a passive fixation task. Our experimental protocol was directly derived from a recent human study (2) to permit a direct comparison between the two primate species. Among the areas defined based on an independent retinotopic mapping experiment, we found that V2, V3, V4 and V3A had significant symmetry selective responses (see figures 2 and 4). We also found symmetry-related activations beyond V4 in a location corresponding to V4A as identified using a probabilistic atlas (see figure 6). Interestingly, response levels to rotation and reflection symmetry in all these areas were very similar (compare the t-scores in figures 2-B and 4-B, see also figure 6-B). We observed parametric responses to rotation symmetry in areas V3, V4 and PITd and higher responses for reflection symmetry around two axes rather than one in V3, V4 and V3A. We further discuss the implications of these results below.

The present results contrast with those reported in the only previous study to have explored symmetry responses in humans and macaques (1). These authors used reflection symmetry patterns defined from random dots. Their results indicated that symmetry responses in macaques were much less robust than in humans, and in fact quite difficult to detect. They concluded that symmetry processing is generally weaker in monkeys than in humans. The results of our study point toward a different conclusion, namely that macaques are quite capable of processing symmetries, at least when they are embedded in regular textures like the ones used here. Indeed, we measured significant responses to reflection and rotation symmetry in a large network of areas which, as we will discuss below, is very consistent with the set of areas found in humans. This was found using BOLD measurements at 3 Teslas with a number of runs in line with the one used in previous studies from our group (30, 16).

BOLD responses to symmetric patterns were consistent in area V2 (t-scores > 3) whereas they were marginal in V1. However, we did not observe significant parametric modulation of V2 responses with rotation symmetry order (experiment 1, figure 3-B) nor stronger responses for reflection symmetry around two axes rather than one (experiment 2, figure 5-B). These results suggest that macaque V2 is capable of a preliminary form of symmetry processing that is less sophisticated than in other areas (see below). The increased sensitivity to image structure in V2 is consistent with a previous study which found that single-cell responses in V2, but not in V1, were sensitive to synthetic stimuli replicating the higher-order statistical dependencies found in natural texture images (18). In humans, V2 has generally not been found to be responsive to symmetry, although Van Meel et al. (11) reported a higher inter-hemispheric connectivity in this area during symmetry perception. Perhaps sensitivity to symmetry arises at an earlier processing stage in macaques than in humans.

Responses to rotation and reflection symmetry were strong in V3. This in line with previous findings in humans (see ref. 2 for rotation symmetry and refs. 1,10 and 11 for reflection symmetry). In our data, V3 was the earliest area in the macaque visual processing stream with parametric responses to rotation symmetry (figure 3-B). Using the same protocol as was used in our first experiment, Kohler et al. (2) also found strong parametric responses to rotation symmetry in human V3, but not V2. In our second experiment, we found that V3 responses were significantly higher for reflection symmetry around two axes (i.e. in the PMM condition) rather than one (figure 5-B). This suggests that macaque V3 is sensitive to the number of reflection symmetry axes as well as order of rotation symmetry. V3 receives inputs from V1 and V2 and projects notably to V4 (19, 20). EEG source localization in humans (2) has suggested that symmetry responses in V3 are unlikely to reflect feedback from higher-level visual areas. It could therefore constitute an important step for the feedforward integration of symmetry and more generally of forms and textures (see ref. 19 for a characterization of V3 responses to higher-order forms). Area V3 is often omitted in the current models of visual processing along the ventral stream (see ref. 21 or 22). Our results suggest that V3 plays an important role in sophisticated form processing typically ascribed to the ventral stream and calls for its incorporation in future models.

Symmetry responses were also strong in area V4 and they share the properties already observed in V3. This is not surprising given that macaque V4 is known to process forms (23) and textures (24). Many commonalities exist between human and monkey V4 (25) and accordingly, our results are in agreement with the fMRI activations observed in humans for rotation (2) and reflection (1,10,11) symmetry in this area. In both species, V3 and V4 could realize an intermediate processing of symmetric patterns before more sophisticated treatments in downstream areas of the ventral pathway. Indeed, human studies consistently reported that areas VO1 (and VO2) and object selective regions like the lateral occipital complex (LOC) play an important role in symmetry perception (26,27). In macaque, our analyses in regions V4A and PITd defined from probabilistic maps (figure 6) showed that area V4A (a potential homologous of human VO1, see ref. 28) had significant responses to both rotation and reflection symmetry (t-scores > 3). PITd, which is strongly activated by shapes and forms (29), as human LOC, had parametric responses to rotation symmetry.

In the dorsal pathway, we found significant symmetry activations in macaque V3A, in agreement with all the previous studies which explored reflection symmetry in its human counterpart (1,10,11). Reflection responses in this area were significantly higher for reflection symmetry around two axes (i.e. in the PMM condition) rather than one. We did not observe parametric responses to rotation symmetry order which is also in line with the human results of Kohler et al. (2). This result suggests an interesting distinction between the ventral and dorsal pathways with only the former having parametric responses to rotation symmetry.

Altogether, our results suggest that the cortical networks that process reflection and rotation symmetry in humans and macaques are rather similar. They call for a deeper exploration of their functional homologies in future studies and open the door to a characterization of the underlying neural mechanisms at the cellular level, notably in areas V2 and V3. These early visual areas represent the first stage of symmetry sensitivity and the transition to a more sophisticated parametric dependence on symmetry order, respectively.

## Materials and methods

### Subjects

Two female rhesus macaques M01 and M02 (age: 17–18 years; weight: 5.30–5,80 kg) were involved in this study. This project was approved by the French Ministry of Research (MP/03/34/10/09) and a local ethics committee (CNREEA code: C2EA – 14). The housing and all the experimental protocols such as surgery, behavioral training, and fMRI recordings (see details on ref. 30) were conducted with respect of the European Union legislation (2010/63/UE) and of the French Ministry of Agriculture guidelines (décret 2013–118). As required and recommended for primate welfare, the two animals were housed together in a social group of 4 individuals into a spacious and enriched enclosure and could thereby develop species-specific behavior such as foraging and congeners delousing.

### Visual stimuli with rotation or reflection symmetry

Our stimuli were defined from previous mathematical works on symmetry based on wallpaper patterns (ref. 31,15). Wallpaper patterns are repetitive 2D patterns that tile the plane. There are 17 unique wallpaper patterns, which cover all planar symmetry groups. Each group is built from a ‘unit lattice’ that is used for tiling the plane without gaps. The “fundamental region” is the smallest repeating region in the wallpaper patterns. Within the unit lattice, multiple rigid transformations are applied to the fundamental region, which give rise to symmetries within the wallpaper group. Each wallpaper group contains a distinct combination of four fundamental symmetries: translations, rotations, reflections, and glide reflections (see ref. (2) for more details). In our experiments, wallpaper patterns were formed by square images of 7 by 7 unit lattices. These square images were subsequently cropped by a circular aperture (11.9° of diameter). In order to characterize the cortical responses to different types of symmetry in macaque, we used here stimuli with either rotation (experiment 1) or reflection (experiment 2) symmetries. The rotation symmetry stimuli were identical to those used in a previous human fMRI study (2) and belonged to four different wallpaper groups: P2, P3, P4 and P6. All four groups contained translation and rotation symmetry but differed in the maximum number of rotations that let the stimuli unchanged. Indeed, rotation symmetry around a point can be defined in terms of its order n, where a rotation by an angle of 360/n does not modify the stimuli. Stimuli from P2, P3, P4 and P6 groups are therefore respectively invariant to rotation of 180, 120, 90 and 60°. Stimuli were generated from a noise texture in which a “fundamental region” was first defined and then repeated and rotated around several points, according to the group’s order of rotation that they belong to (see figure 1-A). The reflection symmetry stimuli were generated using the same procedure but belong to two distinct wallpaper groups, PM and PMM. Both contain reflection and translation symmetry, but while PM has reflection symmetry axes only in one direction, PMM contains axes in two orthogonal directions. We generated versions of PM that had either horizontal or vertical axes of reflection and labelled them PM_v (vertical), and PM_h (horizontal). PMM had reflection axes in both the vertical and horizontal direction (see figure 1-B). For each stimulus exemplar within each wallpaper group (P2, P3, P4, P6, PM_h, PM_v and PMM), we defined a control by applying a 2D-Fourier transform, scrambling the phases of the Fourier coefficients at each frequency and computing the inverse Fourier transform (see figure 1-A). This operation leaves the amplitude spectrum unchanged and therefore preserves the stimulus local properties (luminance, orientation, spatial frequency…). It nonetheless disrupts the global spatial content and controls thereby always degenerate to the simplest wallpaper groups, P1, which contains only translation symmetry.

### MRI recordings

Recordings were performed using a 3 Tesla clinical MR scanner (Philips Achieva) and a dedicated custom 8-channel phased array coil (RapidBiomed) specially designed to fit with the macaque head shape while preserving their field of view.

### Recordings for Individual Templates

Individual structural and functional templates were estimated for our two animals from recording sessions acquired under a slight anaesthesia (Zoletil 100:10mg/kg and Domitor: 0.04 mg/kg) controlled with an MR compatible oximeter. These recordings consisted in four T1-weighted anatomical volumes at high resolution (MPRAGE; repetition time, TR = 10.3 ms; echo time, TE = 4.6 ms, flip angle = 8°; FOV: 155×155 mm; matrix size: 312×192 mm; voxel size = 0.5 × 0.5 × 0.5mm; 192 sagittal slices acquired in an interleaved order) and 300 functional volumes (gradient-echo EPI; TR = 2,000 ms, TE = 30 ms, flip angle = 75°, SENSE factor = 1.6; FOV: 100×100 mm; matrix size: 68×64 mm; voxel size = 1.25 × 1.25 × 1.5mm, 32 axial slices acquired in an interleaved order with a thickness of 1.5 mm and no gap). Anatomical and functional individual templates were derived from those volumes using a procedure that is described in detail in (30).

### Functional recordings

fMRI recordings were conducted on awake behaving animals on a daily basis and lasted about an hour (∼10 runs). The animals were head-fixed, seated in a sphinx position within their primate chair, facing an LCD screen (field of view: 11.9° x 11.9°, resolution: 900 by 900 pixels) at a viewing distance of 1,25 m. They were involved in a passive fixation task while the position of one eye was monitored with an infrared video-based eye-tracker at 500 Hz (Cambridge Research) placed on top of the primate chair. They were water-rewarded during correct fixation (i.e. when their gaze was within a circle of 1° radius around the central fixation point).

For each symmetry condition (P2, P3, P4 and P6 in experiment 1 and PM_h, PM_v and PMM in experiment 2), the main stimuli and their corresponding controls were presented using a block-design (see figure 1-C). Each run consisted of 234s (117 TRs) divided into three identical cycles of 72s (36 TRs) plus an additional baseline of 18s (9 TRs) during which only the fixation point was present. Each cycle started with a baseline of 18s (9 TRs) during which only a gray screen was presented (its luminance was equal to the average luminance in the symmetry stimuli and in their controls). In half of the runs, it was followed by a block of 18s with symmetric stimuli, then by another 18s of baseline and finally by a block of 18s with control stimuli. During a 18s block, a new stimulus appeared every 500ms and therefore there were 36 different stimuli in total. In the other half of the runs, the sequence was reversed and the first baseline of the cycles was followed by a block of control stimuli. Both types of runs were intermixed. The 36 control stimuli in the control blocks corresponded to the 36 stimuli in the symmetric block of the same cycle. The whole experiment (i.e. visual display, eye monitoring and water reward) was controlled using the EventIDE software (Okazolab).

## Data processing

### Individual Templates of Reference

Data collected during the anesthetized sessions (see above) were used to estimate individual functional and anatomical templates. The anatomical template was obtained by realigning and averaging the 4 T1-weighted (MPRAGE) volumes. It was then aligned to the MNI space of the 112RM-SL template (see ref. 32, 33). Cortical surface reconstructions were performed using the CARET software (34).The functional template was obtained by realigning and averaging the 300 functional (GE-EPI) volumes. It was aligned with the anatomical template and spatial normalization parameters (affine and non-rigid) between the functional and anatomical templates were determined based on the gray matter maps of both templates. For group analyses, the same operation was performed to register each individual anatomical template to the F99 template available in the CARET software (35).

### Preprocessing of the Functional Data

To minimize the influence of eye movements on our results, only runs with high fixation rate (> 85%) were considered for further analyses. For our two monkeys (M01 and M02), we respectively collected 16 and 25 of such runs for each rotation symmetry groups (i.e. P2, P3, P4 and P6) in the first experiment. We also collected 18 (M01) and 16 (M02) of such runs for each reflection symmetry group (PM_h, PM_v and PMM) in the second experiment. For each experiment, the different symmetry conditions were interleaved between runs. The four first volumes of each run were removed to account for signal stabilization. The remaining 113 volumes were then realigned and corrected for slice timing before being co-registered to the functional template first and finally to the anatomical template. Images were then smoothed using a spatial Gaussian kernel with a FWHM of 2 mm^3^.

### General linear model (GLM)

Voxel-wise statistics were computed by fitting a general linear model (GLM) to the BOLD signal. The model contained 3 main regressors, representing the 3 experimental conditions: symmetry (rotation for experiment 1 and reflection for experiment 2), control, and blank periods. These regressors were convolved with the hemodynamic response function (HRF) estimated from each of the two monkeys (see details in ref.16). To eliminate noise in our recordings, we performed a principal component analysis on voxels located outside the brain (see ref. 36). Time-courses in those voxels mostly reflect artifacts caused by movement of the animals and should be independent of our experimental design. For each run, we determined the number of principal components that were necessary to explain 80% of the variance in these voxels and used the corresponding principal vectors as regressors of non interest in our model.

The beta weights obtained from the GLM were subsequently used to perform univariate analyses (t-scores) at the whole brain level. These analyses were performed on the preprocessed EPI data and both the beta weights and the associated t-scores were then projected onto the high-resolution volumes of our two animals. They were also projected on the individual cortical surfaces (see figures 2-A, 3-A, 4-A and 5-A) and on the cortical surface of the F99 template (see supplementary figure 2) using the Caret software (34). All the preprocessing and GLM analyses were executed using the Matlab and SPM12 softwares.

### Analyses in retinotopically defined ROIs

For our two animals, we used a population receptive field (pRF) analysis to define visual areas V1, V2, V3, V3A and V4 on the individual cortical surfaces from the data collected during an independent wide-field retinotopic experiment (see ref. (16) and (17)). The same data were used to define the MT and PIP clusters and their satellite sub-regions (V4t, MT, MSTv and FST for the MT cluster and CIP1, CIP2, PIP1 and PIP2 for the PIP cluster). Because the field of view used for the symmetry experiment (11.9° x 11.9°, see above) was much smaller than during our retinotopic mapping procedure, we restricted these areas and clusters by keeping only the cortical nodes with significant visual responses during the symmetry experiment (i.e. nodes with t-scores > 3 for the contrast between vision and baseline). This operation prevents from including nodes whose receptive fields are outside the stimulated area in the analyses. Within each of these restricted retinotopic ROIs and clusters, we computed the average beta values for each experimental run and each monkey (data from both hemispheres were combined). These average values were used to perform ROI-level statistical analyses: for each experiment, we estimated the t-scores between the betas obtained in all the symmetry conditions (P2, P3, P4 and P6 for experiment 1, PM_h, PM_v and PMM for experiment 2) and those obtained in the associated control conditions. Note that in the corresponding figures (figure 2-B and figure 4-B), we show these data averaged across the two monkeys. We also provide t-scores in the subparts of the ROIs corresponding to the left and right hemispheres of M01 and M02 to demonstrate their very good correspondence and hence illustrate the robustness and reproducibility of our data.

### Comparisons between responses to different symmetry conditions

In humans, some visual areas have rotation symmetry responses that are proportional to the symmetry order (2). To test whether such properties also exist in macaque, we computed for all our ROIs the percentages of signal change (PSCs) between each rotation symmetry condition (P2, P3, P4 and P6) and their respective controls (see figure 3-B). We subsequently performed linear regressions between these PSCs and symmetry order for each monkey using the lm1 package in R (RCore Team, 2014). We considered that a ROI had rotation symmetry responses significantly proportional to the symmetry order only when we found a significant (p-value < 0.05) linear relationship for both monkeys. We chose this conservative criterion to avoid false positives.

PSCs were also used to compare the responses to the different reflection symmetry conditions (experiment 2). These comparisons were performed using two-tailed non-parametric permutation analyses. Here as well, we considered that responses in an ROI were significantly stronger for one condition than for another when significant statistical differences (p-value < 0.05) were found in both the two monkeys.

### Analyses in ROIs defined from a probabilistic atlas

In order to characterize activations beyond retinotopic area V4, we used the probabilistic maps of area V4A, dorsal occipitotemporal area (OTd) and ventral and dorsal posterior inferotemporal areas (PITv and PITd) provided by Janssens and Vanduffel (see ref. 37) in the CARET software. For each of these areas, we selected the nodes of the F99 template with a probability score above 0.8 and projected the associated binary maps on the inflated reconstructions of the left and right cortical hemispheres of our two monkeys (M01 and M02).

## Acknowledgments

This work was supported by the Agence Nationale de la Recherche Jeunes Chercheuses et Jeunes Chercheurs (Grants ANR-16-CE37-0002-01, 3D3M) awarded to BRC. We thank the Inserm/UPS UMR1214 Technical Platform for the MRI acquisitions. We also thank the Animalliance staff for their help with monkey welfare. PJK acknowledges funding support from the Canada First Research Excellence Fund and the National Sciences and Engineering Research Council of Canada.

## References

1. Y. Sasaki, W. Vanduffel, T. Knutsen, C. Tyler, R. Tootell, Symmetry activates extrastriate visual cortex in human and nonhuman primates. Proc. Natl. Acad. Sci. U. S. A. 102, 3159–3163 (2005).

2. P. J. Kohler, A. Clarke, A. Yakovleva, Y. Liu, A. M. Norcia, Representation of Maximally Regular Textures in Human Visual Cortex. J. Neurosci. Off. J. Soc. Neurosci. 36, 714–729 (2016).

3. A. P. Lightman, The accidental universe: the world you thought you knew, 1. American ed (Pantheon Books, 2013).

4. J. D. Delius, B. Nowak, Visual symmetry recognition by pigeons. Psychol. Res. 44, 199–212 (1982).

5. J. Merry, M. R. Morris, Preference for symmetry in swordtail fish. Anim. Behav. (2001) https://doi.org/10.1006/anbe.2000.1589.

6. M. Giurfa, B. Eichmann, R. Menzel, Symmetry perception in an insect. Nature 382, 458–461 (1996).

7. A.P. Møller, Female swallow preference for symmetrical male sexual ornaments. Nature 357, 238–240 (1992).

8. J. Wagemans, Characteristics and models of human symmetry detection. Trends Cogn. Sci. 1, 346–352 (1997).

9. T. Jacobsen, L. Höfel, Descriptive and evaluative judgment processes: Behavioral and electrophysiological indices of processing symmetry and aesthetics. Cogn. Affect. Behav. Neurosci. 3, 289–299 (2003).

10. B. D. Keefe, et al., Emergence of symmetry selectivity in the visual areas of the human brain: fMRI responses to symmetry presented in both frontoparallel and slanted planes. Hum. Brain Mapp. 39, 3813–3826 (2018).

11. C. Van Meel, A. Baeck, C. R. Gillebert, J. Wagemans, H. P. Op de Beeck, The representation of symmetry in multi-voxel response patterns and functional connectivity throughout the ventral visual stream. NeuroImage 191, 216–224 (2019).

12. C. Waitt, A. C. Little, Preferences for Symmetry in Conspecific Facial Shape Among Macaca mulatta. Int. J. Primatol. 27, 133–145 (2006).

13. D. J. Felleman, D. C. Van Essen, Distributed hierarchical processing in the primate cerebral cortex. Cereb. Cortex N. Y. N 1991 1, 1–47 (1991).

14. G. A. Orban, D. Van Essen, W. Vanduffel, Comparative mapping of higher visual areas in monkeys and humans. Trends Cogn. Sci. 8, 315–324 (2004).

15. Y. Liu, H. Hel-Or, C. S. Kaplan, L. Van Gool, “Human Visual Perception of Two-Dimensional Symmetry” (PsyArXiv, 2010) https://doi.org/10.31234/osf.io/x7uvf (March 1, 2021).

16. Y. Héjja-Brichard, S. Rima, E. Rapha, J.-B. Durand, B. R. Cottereau, Stereomotion Processing in the Nonhuman Primate Brain. Cereb. Cortex N. Y. N 1991 30, 4528–4543 (2020).

17. S. Rima, B. R. Cottereau, Y. Héjja-Brichard, Y. Trotter, J.-B. Durand, Wide-field retinotopy reveals a new visuotopic cluster in macaque posterior parietal cortex. Brain Struct. Funct. 225, 2447–2461 (2020).

18. J. Freeman, C. M. Ziemba, D. J. Heeger, E. P. Simoncelli, J. A. Movshon, A functional and perceptual signature of the second visual area in primates. Nat. Neurosci. 16, 974–981 (2013).

19. D. J. Felleman, A. Burkhalter, D. C. Van Essen, Cortical connections of areas V3 and VP of macaque monkey extrastriate visual cortex. J. Comp. Neurol. 379, 21–47 (1997).

20. K. R. Gegenfurtner, D. C. Kiper, J. B. Levitt, Functional properties of neurons in macaque area V3. J. Neurophysiol. 77, 1906–1923 (1997).

21. M. Riesenhuber, T. Poggio, Models of object recognition. Nat. Neurosci. 3, 1199–1204 (2000).

22. J. J. DiCarlo, D. D. Cox, Untangling invariant object recognition. Trends Cogn. Sci. 11, 333–341 (2007).

23. R. Desimone, S. J. Schein, Visual properties of neurons in area V4 of the macaque: sensitivity to stimulus form. J. Neurophysiol. 57, 835–868 (1987).

24. G. Okazawa, S. Tajima, H. Komatsu, Image statistics underlying natural texture selectivity of neurons in macaque V4. Proc. Natl. Acad. Sci. 112, E351–E360 (2015).

25. A. W. Roe, et al., Toward a Unified Theory of Visual Area V4. Neuron 74, 12–29 (2012).

26. P. J. Kohler, B. R. Cottereau, A. M. Norcia, Dynamics of perceptual decisions about symmetry in visual cortex. NeuroImage 167, 316–330 (2018).

27. S. Bona, A. Herbert, C. Toneatto, J. Silvanto, Z. Cattaneo, The causal role of the lateral occipital complex in visual mirror symmetry detection and grouping: an fMRI-guided TMS study. Cortex J. Devoted Study Nerv. Syst. Behav. 51, 46–55 (2014).

28. M. J. Arcaro, M. S. Livingstone, Retinotopic Organization of Scene Areas in Macaque Inferior Temporal Cortex. J. Neurosci. 37, 7373–7389 (2017).

29. H. Kolster, T. Janssens, G. A. Orban, W. Vanduffel, The Retinotopic Organization of Macaque Occipitotemporal Cortex Anterior to V4 and Caudoventral to the Middle Temporal (MT) Cluster. J. Neurosci. 34, 10168–10191 (2014).

30. B. R. Cottereau, et al., Processing of Egomotion-Consistent Optic Flow in the Rhesus Macaque Cortex. Cereb. Cortex N. Y. NY 27, 330–343 (2017).

31. E. S. Fedorov, The Symmetry of Regular Systems of Figures (in Russian). Proc. Imp. StPetersburg Mineral. Soc., 1–146 (1891).

32. D. G. McLaren, et al., A population-average MRI-based atlas collection of the rhesus macaque. NeuroImage 45, 52–59 (2009).

33. D. G. McLaren, K. J. Kosmatka, E. K. Kastman, B. B. Bendlin, S. C. Johnson, Rhesus macaque brain morphometry: a methodological comparison of voxel-wise approaches. Methods San Diego Calif 50, 157–165 (2010).

34. D. C. Van Essen, et al., Mapping visual cortex in monkeys and humans using surface-based atlases. Vision Res. 41, 1359–1378 (2001).

35. D. C. Van Essen, Surface-based atlases of cerebellar cortex in the human, macaque, and mouse. Ann. N. Y. Acad. Sci. 978, 468–479 (2002).

36. W. Vanduffel, R. Farivar, “Functional MRI of Awake Behaving Macaques Using Standard Equipment” in Advanced Brain Neuroimaging Topics in Health and Disease - Methods and Applications, T. D. Papageorgiou, G. I. Christopoulos, S. M. Smirnakis, Eds. (InTech, 2014) https://doi.org/10.5772/58281 (March 2, 2021).

37. T. Janssens, Q. Zhu, I. D. Popivanov, W. Vanduffel, Probabilistic and single-subject retinotopic maps reveal the topographic organization of face patches in the macaque cortex. J. Neurosci. Off. J. Soc. Neurosci. 34, 10156–10167 (2014).

