## Supplementary_Information for "Symmetry processing in the macaque visual cortex"

**This PDF file includes:**

**Supplementary figures S1 to S6**

**Supplementary table S1**

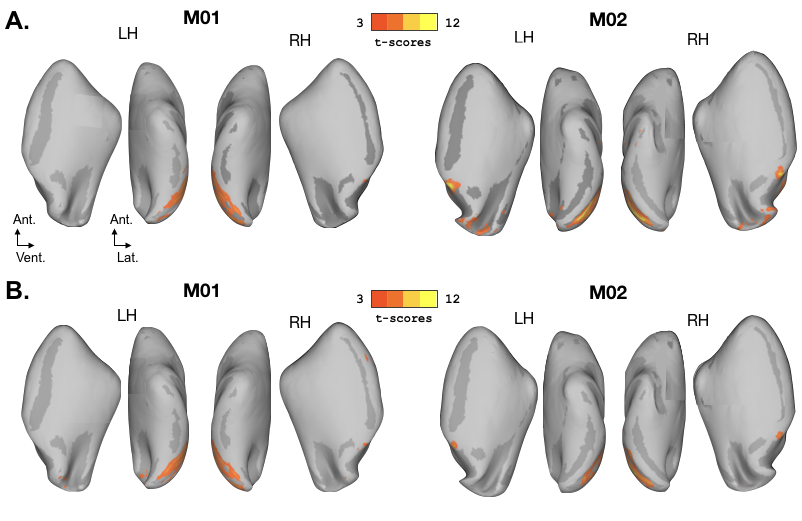

**Supplementary figure 1:** Medial and ventral views of the activations obtained for rotation (panel A) and reflection (panel B) symmetries. Data were thresholded at p-value < 10^−3^ (uncorrected). Ant.: anterior. Vent.: ventral. Lat.: lateral. LH: left hemisphere. RH: right hemisphere. See figure 2-A and 4-A for more details.

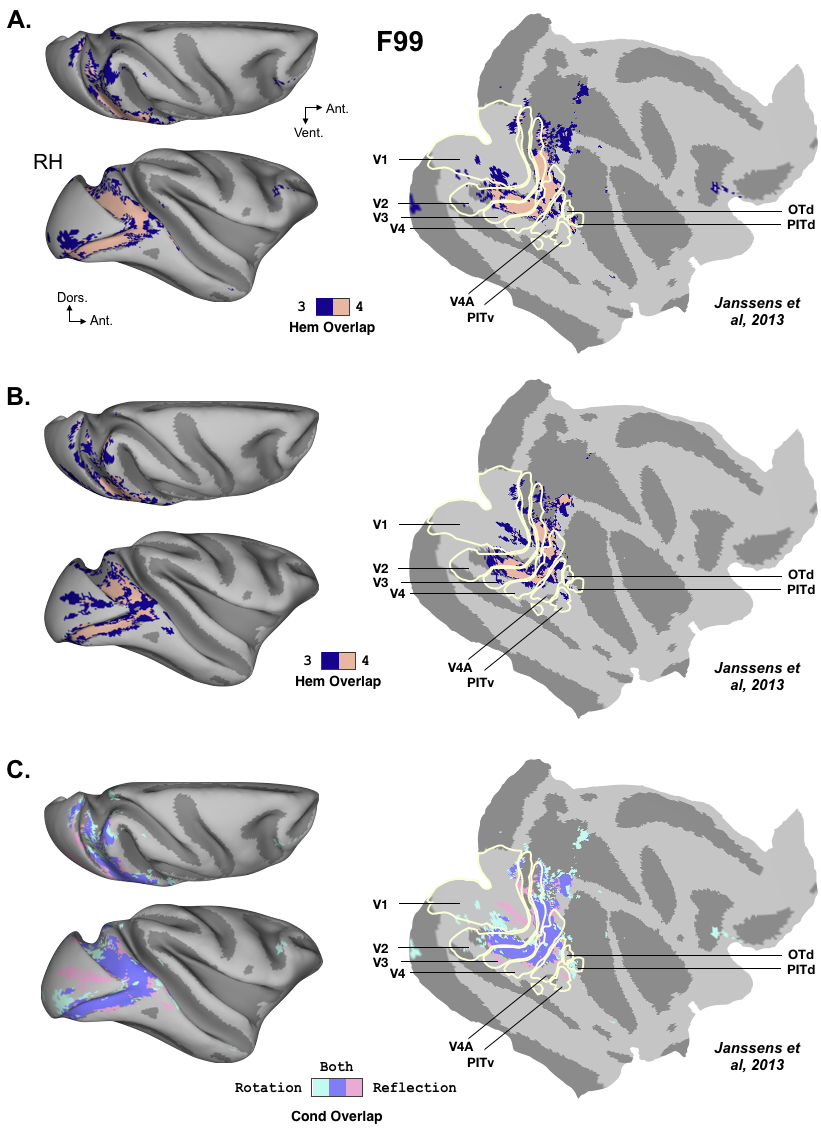

**Supplementary figure 2:** Overlap maps between the activations obtained in the two animals. Data were projected on the right hemisphere of the F99 template. ROI borders (in light yellow) were defined from the probability maps described by Janssens et al. (37). A) Overlap maps for the contrast between the rotation symmetry conditions and their controls (see figure 2-A). These maps provide cortical nodes where significant effects (p-value < 10^-3^, uncorrected) were found in 3 (dark blue) or 4 (salmon) hemispheres. B) Overlap maps for the contrast between the reflection symmetry conditions and their controls (see figure 4-A). C) Overlap maps between the results of the first (rotation symmetry) and second (reflection symmetry) experiments. Nodes where significant effects were found in at least 3 hemispheres for the first, second and both experiments are respectively shown in cyan, pink and purple.

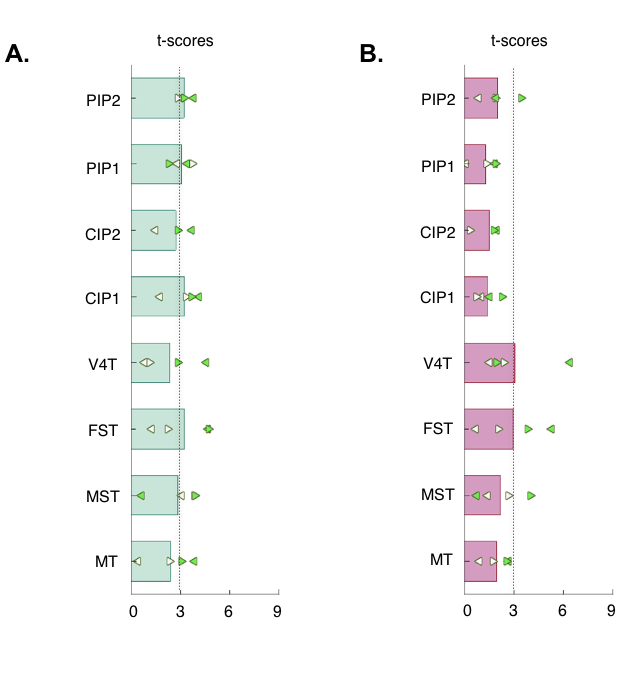

**Supplementary figure 3:** Symmetry responses within the satellite areas of the MT (V4t, MT, FST and MSTv) and PIP (CIP1/2 and PIP1/2) clusters. A) T-scores for the rotation symmetry versus control conditions (experiment 1). See figure 2-B for more details. B) T-scores for the rotation symmetry versus control conditions (experiment 1). See figure 4-B for more details.

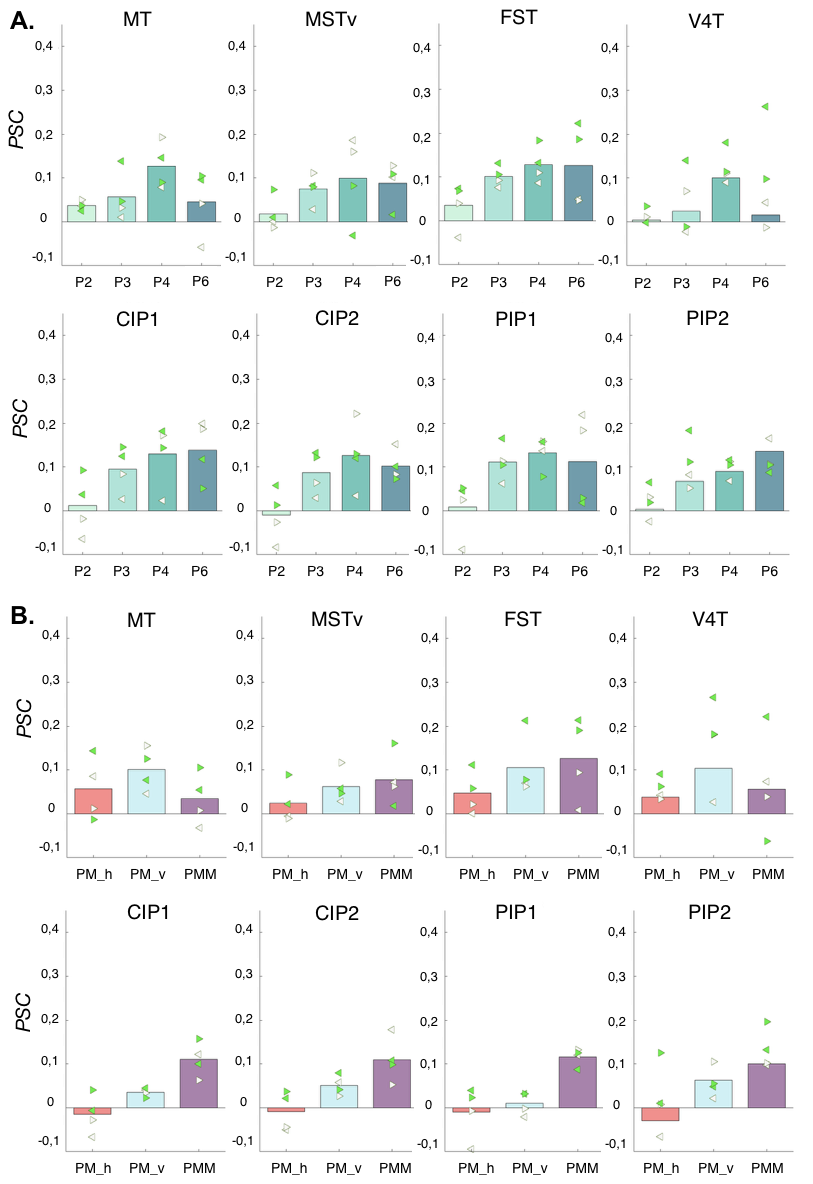

**Supplementary figure 4:** Symmetry effects within the satellite areas of the MT (V4t, MT, FST and MSTv) and PIP (CIP1/2 and PIP1/2) clusters. A) Percentages of signal changes (PSCs) obtained for each of the rotation symmetry conditions (P2, P3, P4 and P6) versus their respective controls. See figure 3-B for more details. B) Percentages of signal changes (PSCs) obtained for each of the reflection symmetry conditions (PM_v, PM_h and PMM) versus their respective controls. See figure 5-B for more details.

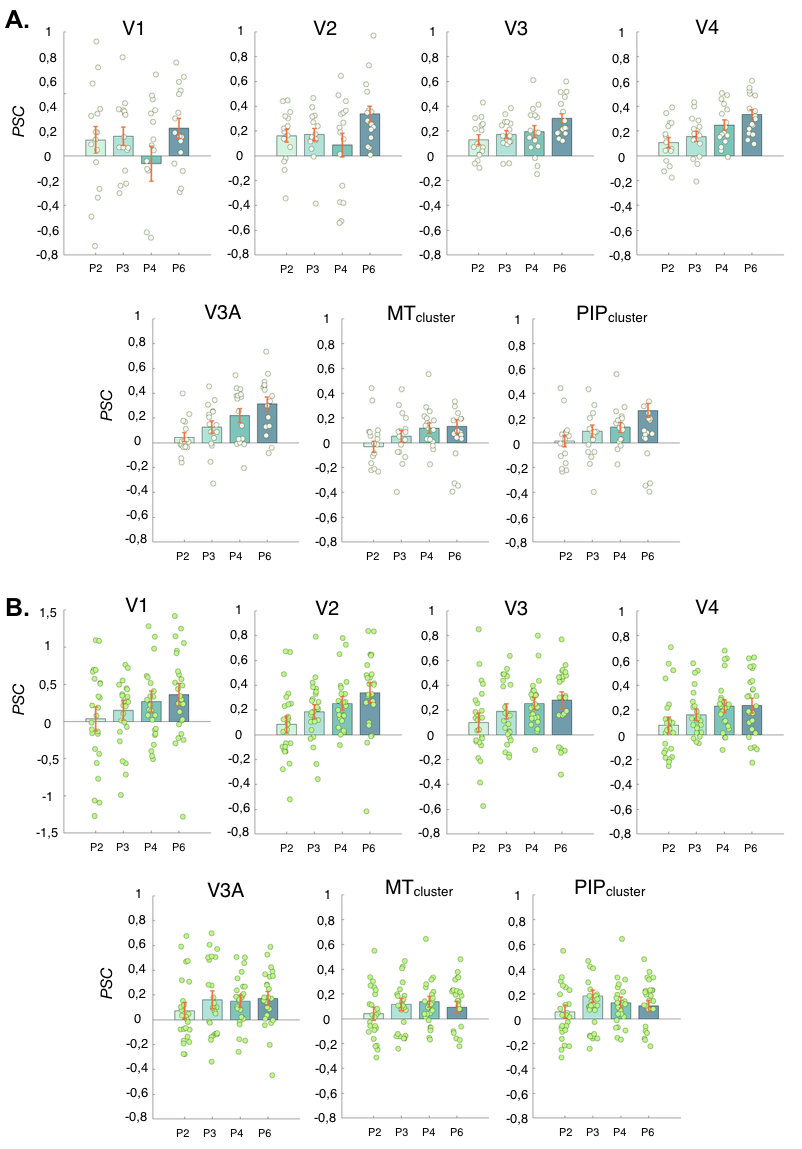

**Supplementary figure 5:** Effects of rotation symmetry order on BOLD responses in monkey M01 (A) and M02 (B). Percentages of signal changes (PSCs) between the responses to the rotation symmetry conditions (P2, P3, P4 and P6) and those to their respective controls. Data are shown in retinotopic areas (V1, V2, V3, V3A and V4) and in the MT and PIP clusters. Each circular data point corresponds to the PSC measured in one run. For each rotation symmetry order, the red bar gives the standard errors of the mean across runs (s.e.m., n = 16 for M01 and n = 25 for M02).

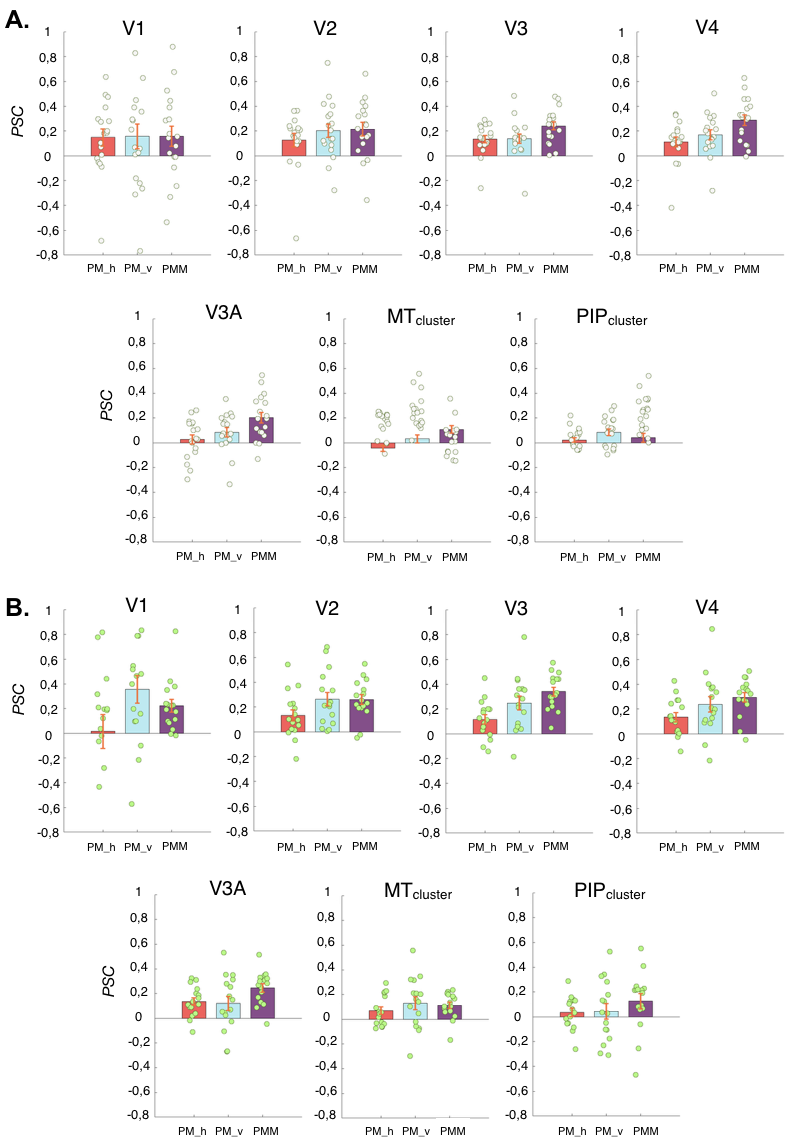

**Supplementary figure 6:** Effects of the different reflection symmetry conditions on BOLD responses in monkey M01 (A) and M02 (B). Percentages of signal changes (PSCs) obtained for each of the reflection symmetry conditions (PM_h, PM_v and PMM) versus their respective controls. Data are shown in retinotopic areas (V1, V2, V3, V3A and V4) and in the MT and PIP clusters. Each circular data point corresponds to the PSC measured in one run. For each condition, the red bar gives the standard errors of the mean across runs (s.e.m., n = 18 for M01 and n = 16 for M02).

| ROI | Subject | Equation | Variance explained (%) |
| --- | --- | --- | --- |
| V3 | M01 | PSC = 0,042x + 0,039 | 0,981 |
|  | M02 | PSC = 0,043x + 0,043 | 0,835 |
| V4 | M01 | PSC = 0,057x - 0,006 | 0,975 |
|  | M02 | PSC = 0,037x + 0,035 | 0,761 |
| PITd | M01 | PSC = 0,029x -0,020 | 0,992 |
|  | M02 | PSC = 0,048x - 0,069 | 0,855 |

**Supplementary table 1:** Linear equations between PSC and rotation symmetry order (x) in visual ROIs for which we found a significant linear effect (p-value < 0.05) in both monkeys. None of the associated coefficients had a confidence interval that included 0. We also provide the variances of the average PSC explained by the models (rightward column).
